# Long-horizon associative learning explains human sensitivity to statistical and network structures in auditory sequences

**DOI:** 10.1101/2024.01.16.575814

**Authors:** Lucas Benjamin, Mathias Sablé-Meyer, Ana Fló, Ghislaine Dehaene-Lambertz, Fosca Al Roumi

## Abstract

Networks are a useful mathematical tool for capturing the complexity of the world. In a previous behavioral study, we showed that human adults were sensitive to the high-level network structure underlying auditory sequences, even when presented with incomplete information. Their performance was best explained by a mathematical model compatible with associative learning principles, based on the integration of the transition probabilities between adjacent and non-adjacent elements with a memory decay. In the present study, we explored the neural correlates of this hypothesis via magnetoencephalography (MEG). Participants passively listened to sequences of tones organized in a sparse community network structure comprising two communities. An early difference (~150 ms) was observed in the brain responses to tone transitions with similar transition probability but occurring either within or between communities. This result implies a rapid and automatic encoding of the sequence structure. Using time-resolved decoding, we estimated the duration and overlap of the representation of each tone. The decoding performance exhibited exponential decay, resulting in a significant overlap between the representations of successive tones. Based on this extended decay profile, we estimated a long-horizon associative learning novelty index for each transition and found a correlation of this measure with the MEG signal. Overall, our study sheds light on the neural mechanisms underlying human sensitivity to network structures and highlights the potential role of Hebbian-like mechanisms in supporting learning at various temporal scales.

## Introduction

Understanding the structure of the input sequences we encounter is fundamental for developing a comprehensive mental model of our environment ***(Dehaene et al., 2022, 2015)***. The capacity to detect first-order relationships between successive events (i.e., transition probabilities) and its limits have been extensively studied in humans at the behavioral and neural levels ***(Benjamin et al., 2021, 2023b, 2024; Fló et al., 2022; Henin et al., 2021; Maheu et al., 2019; Saffran et al., 1996)*** as well as in non-humans animals ***(Boros et al., 2021; James et al., 2020; Toro and Trobalón, 2005)***. Higher-order statistical relations between elements of a sequence are also detected by human adults and children ***(Karuza et al., 2019, 2019; Lynn et al., 2020; Mark et al., 2020; Schapiro et al., 2013)***, but only a limited number of neuroimaging studies have explored possible neural correlates of this learning ***(Ren et al., 2022; Schapiro et al., 2016; Stiso et al., 2022)***. Therefore, we still do not know if a single mechanism can adequately explain both first order (local transitions) and network structure learning or if these computations require distinct cognitive and brain processes.

To bridge the gap between local statistical and network-level learning studies, we previously proposed the *sparse community paradigm*, which allows to simultaneously characterize these aspects on auditory sequences ***(Benjamin et al., 2023a)***. Building upon the community paradigm introduced by ***Schapiro et al (2013)***, we created a network consisting of two densely but incompletely connected clusters (called communities) of six elements each. Each element is connected with four other elements, and the two clusters are linked by only two edges (links). A learning sequence is created by randomly drawing the next tone from the four possibilities, creating a random walk in the network with a uniform transition probability (TP) between successive tones (fig 1A). After exposure to such a sequence, participants were asked to judge their familiarity with various pairs of tones that 1) had or had not been presented during learning to test local TP learning, and 2) did or did not belong to the same community to test learning of the higher-level structure ***(Benjamin et al., 2023a)***. Interestingly, participants judged new transitions they had never heard as highly familiar if they were between tones belonging to the same community. This *completion* effect demonstrated that they generalized the community structure to missing transitions. Conversely, they judged transitions between communities to be less familiar than within communities despite the absence of any difference in local transition probability during learning. This *pruning* effect translates into a decrease in subjective familiarity with tone pairs that switch from one community to the other despite similar transition probabilities between tones. Among the various models proposed in the statistical and network learning literature, an associative learning approach (the free energy minimization model - FEMM ***(Lynn et al., 2020)***), conceptually related to the successor representation, provided the best fit to participants’ behavior. According to this model, participants did not solely compute adjacent transition probabilities but a linear sum of transition probabilities at all orders (adjacent, first-order non-adjacent, second-order non-adjacent and so on), weighted by a decreasing exponential factor. This model explains how both local transitions and network structures are perceived, and successfully accounts for behavioral results across different network types, including community, sparse community, ring, and lattice networks ***(Benjamin et al., 2023a; Lynn et al., 2020)***, as well as results concerning local statistical learning. FEMM appears to be a good candidate for a unifying framework of sequence learning.

**Figure 1:**
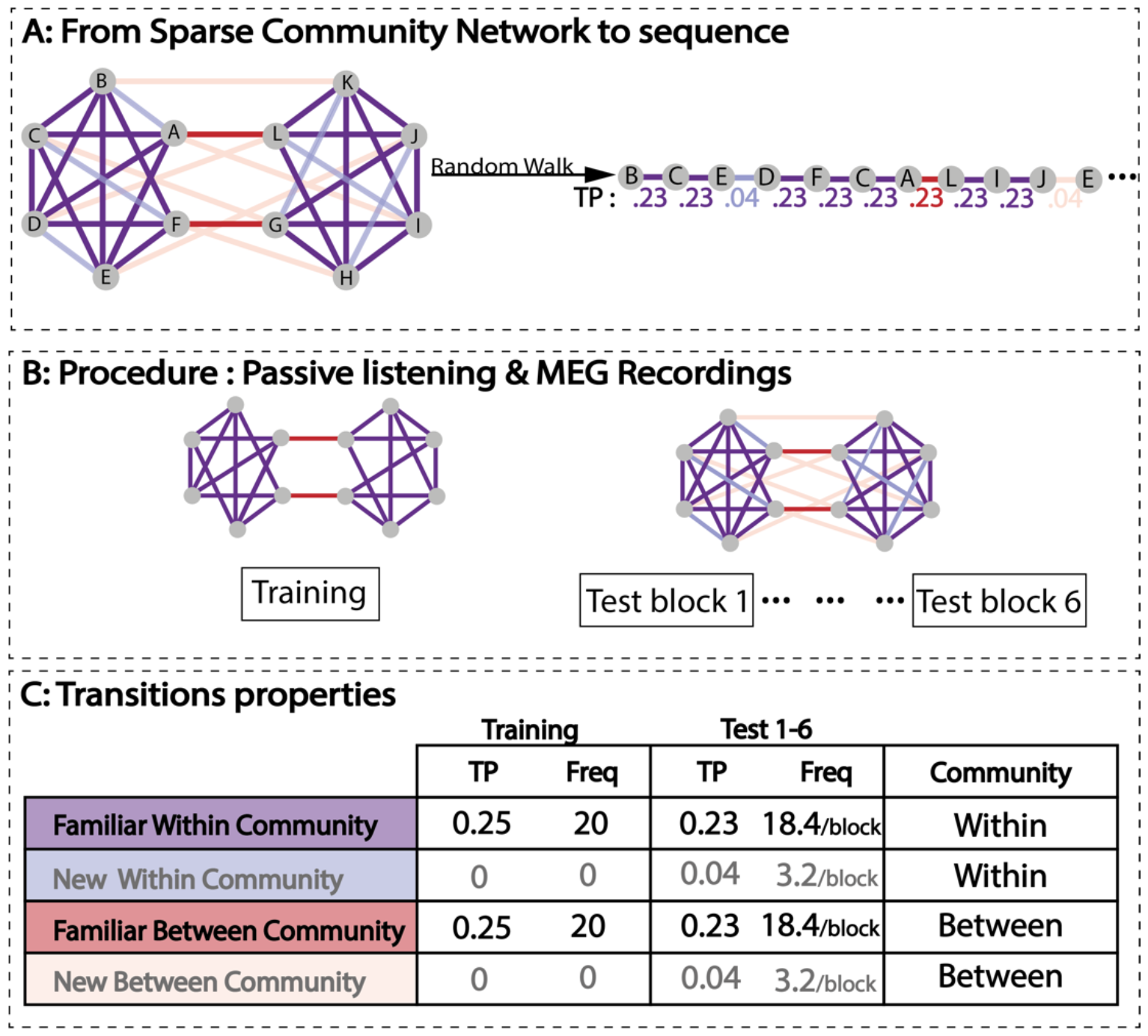
Design and procedure. A) Example of a sparse community network for one participant. All community networks are similar in terms of properties, but New Within and New Between transitions are randomly drawn for each participant. Purple lines correspond to Familiar Within-community transitions, red lines to Familiar Between community transitions, and blue and pink lines correspond respectively to New Within and New Between transitions. We can derive a sequence by performing a random walk into this network. Here we display an example of a test sequence derived from this structure. B) Experimental procedure. First, participants passively listened to a sequence from a sparse community network, in which each TP between tones was 25% (Training). Then they were presented with six 960-items test blocks obtained from the community structure graph comprising New Within and Between community transitions with low transition probabilities of 4% (light blue and pink colors on the graph). C) Table summarizing the local and community properties for the transitions for each condition. Each single Familiar transition is, on average, presented 18.4 times/block (20 in the Training sequence) and, therefore, has a probability of 23% to be observed (25% in the training sequence) irrespectively of staying within or switching between communities. New transitions have a probability of 4% in the test blocks, which implies that each single New Transition is heard 3.2 times/block on average.

However, a common model is insufficient to postulate a common implementation ***(Marr, 1982)*** and there is still no consensus on how the brain implements these computations. On the one hand, sensitivity to network structure is often described as a conscious abstraction of the structure involving top-down attention processes with late brain signatures ***(Ren et al., 2022)*** typically in the prefrontal cortex ***(Stiso et al., 2022)***. On the other hand, we previously postulated that low-level associative learning ***(Benjamin et al., 2023a; see also Endress, 2010; Endress and Johnson, 2021; Schapiro et al., 2017)*** was sufficient for both local and higher-order learning. To disentangle those two hypotheses, we tested here whether passive exposure to a rapid auditory sequence could lead to successful learning of its network structure. We thus exposed participants to fast sequence of tones following the sparse community design while recording their brain activity with MEG.

## Materials and methods

### Stimuli and procedure

We generated twelve tones of 50ms duration, logarithmically distributed from 300 to 1800 Hz. For each participant, the twelve tones were randomly assigned to the twelve nodes of the sparse community network (see fig 1 for a complete description of the network structure). The sparse community network comprised two communities (i.e., clusters) made of six nodes, densely connected to each other but poorly connected to the nodes of the other community. Crucially, in the *sparse* community design, some connections between nodes belonging to the same community are missing. Specifically, for each participant, we randomly removed twelve transitions (6 per community, one per node). After a training block with this incomplete graph, new transitions were added at a low frequency (4%) for the following test blocks. We refer to these transitions as *New* and those presented during training as *Familiar*. The *New* transitions were critically within and between the communities (which we refer to as *Within* vs *Between*). *New Within* corresponds to the twelve ‘missing’ transitions randomly removed from the network in the training block. To balance, we randomly selected 12 *New Between* transitions (one per node) that violated the community clustering property. As a result, the transition probabilities between tones during the training block were flat: TP = 25%, while during the test blocks, the *Familiar* transitions had TP = 23% and a frequency of 18.4/block. The 12 *New Within* and 12 *New Between* community transitions had TP = 4% and a frequency of 3.2/block. The *New Within* and *New Between* transitions were randomly drawn for each subject to add variability to the network structure. Fig 1A shows an example of one structure and the associated sequence used for one participant.

We then performed random walks in the participant’s sparse community graph to derive one 960 items-long training sequence and six 960 items-long test sequences (one sequence corresponds to one block) with 200ms inter stimulus interval (ISI) between each tone (fig 1A). The first block of 960 items comprised only *Familiar Within* and *Familiar Between* transitions (training block, TP = 25% each). For the next six blocks (Test 1-6), we introduced infrequent *New Within* and *New Between* transitions (TP = 4% each). All *Familiar* transitions, independently of whether they were *Within* or *Between* communities, had the same TPs and appeared with the same frequency (TP = 23% each). However, the graph structure entails that the participants heard in total fewer between community transitions than within community transitions (there are 32 *Familiar Within* and 4 *Familiar Between* community transitions during training, completed by *12 New Within* and *12 New Between* community transitions during test).

Crucially, the experiment was completely passive, and participants were unaware of the structure of the auditory sequence. They were only instructed to pay attention to the sequence of tones and to stay still while looking at a fixation cross displayed at the center of the screen to avoid noise from eye movements. The experiment lasted around 45 minutes, and a small break inside the MEG was possible between each block.

### Participants

29 healthy adults came to the lab and 23 recordings (16 females, mean age = 26.58, sd = 6.1) were kept for the analyses (4 subjects were rejected due to MEG malfunction, 1 due to experimenter error during recording, and 1 scan was aborted due to subject agitation). All participants gave written informed consent prior to enrollment and received 90€ as compensation. This experiment was approved by the national ethical committee (CPP Ile-de-France III).

### MEG recordings and preprocessing

Participants performed the tasks while sitting inside an electromagnetically shielded room. The magnetic component of their brain activity was recorded with a 306-channel, whole-head MEG by Elekta Neuromag® (Helsinki, Finland). The MEG helmet is composed of 102 triplets, each comprising one magnetometer and two orthogonal planar gradiometers. Brain signal was acquired at a sampling rate of 1000 Hz with a hardware high-pass filter at 0.03Hz. The data were then resampled at 250Hz to reduce computational load. Eye movements were monitored with vertical and horizontal EOGs and heartbeats with ECGs. Subjects’ head position inside the helmet was measured for realignment at the beginning of each run with an isotrack Polhemus Inc. system from the location of four coils placed over the frontal and mastoids.

MEG signal was then preprocessed using MNE python pipeline with classical steps following recommendations from ***(Jas et al., 2018; Niso et al., 2018)***. We first applied Maxfilter algorithm to remove ambient noise, and signal was band-pass filtered ([0.1-30] Hz). Eye movements and heartbeats were identified and removed using PCA components’ correlation with EOG and ECG measures.

To decode if a transition was within or between community, data was epoched from 100 ms before to 300ms after tone onset. To determine how sustained was the neural representation of each tone across time, we segmented the data in 2.6 seconds long epochs, from 100 ms before to 2500 ms after tone onset. Bad data, channels, and epochs were detected and removed with autoreject toolbox ***(Jas et al., 2017)***.

### Within vs Between Decoding analysis

To examine whether the brain encoded the community structure, we trained a logistic regression decoder to predict whether the transition that just occurred stayed *Within* a community (*Familiar Within* and *New Within*) or switched *Between* communities (*Familiar Between* and *New Between*). The decoder was trained on the short epochs ([-0.1, 0.3]) slightly smoothed using a sliding window (± 20ms) to enhance the signal-to-noise ratio. We used 3 folds cross-validation process: the decoder was trained on 2/3 of the data and tested on the remaining third of the trials. The procedure was repeated three times, corresponding to the 3 cross-validation folds. Each transition had the same frequency but *Within* transitions were more numerous than *Between* transitions, resulting in a larger total number of epochs for *Within* condition. We thus used the area under the ROC curve as a metric of success (ROC AUC) since it is not sensitive to such imbalance. This analysis was conducted for each time-point of the epochs (fig 2). We also computed the decoding performance when the decoder was trained at time t and tested at time t’, to reveal the generalization across time (GAT) of the decoder, and thus the stability of the mental representation (fig 2). By design, the diagonal of the GAT matrix corresponds to the previously described time-by-time decoding performance.

**Figure 2:**
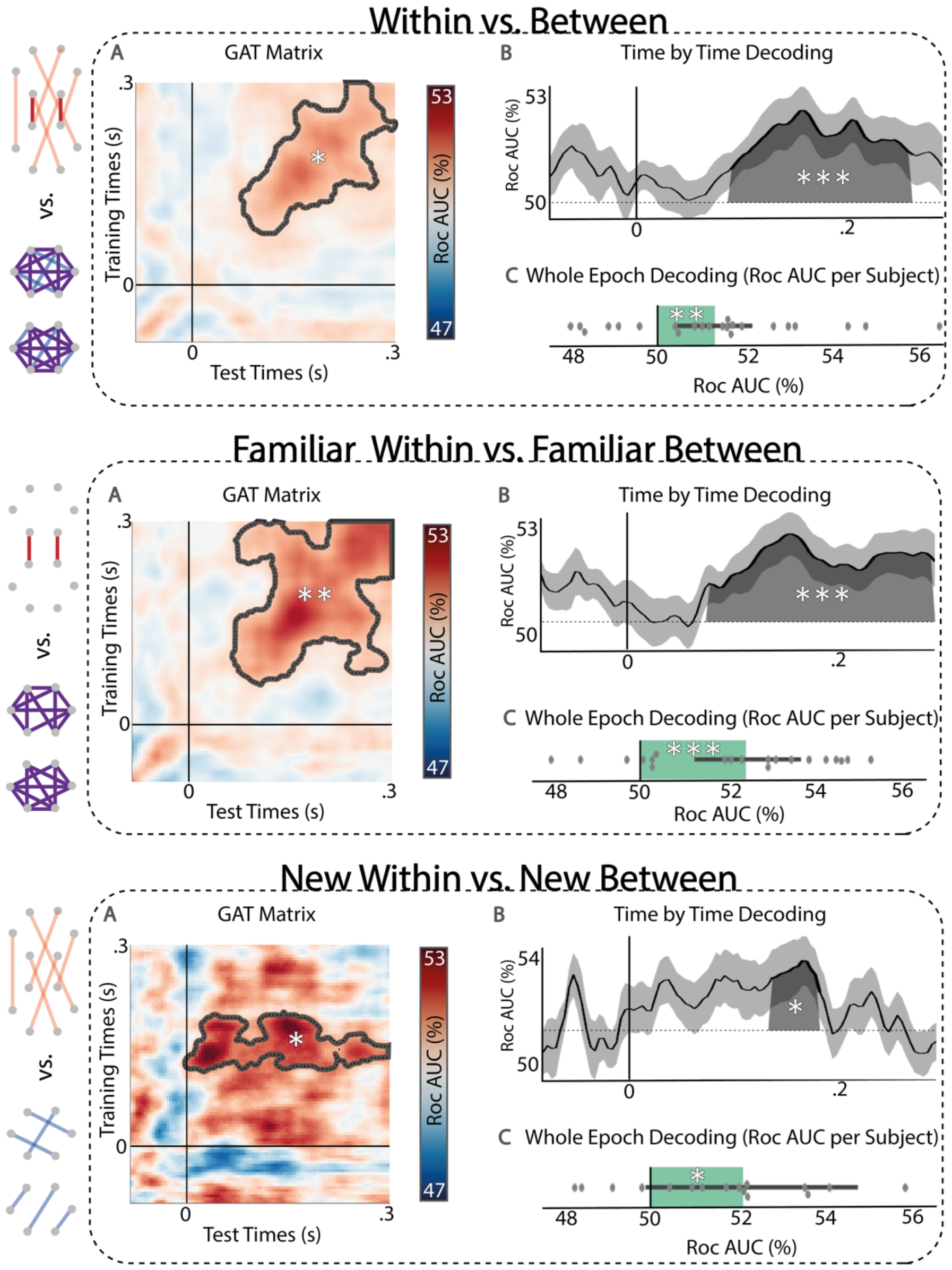
Within vs. Between community decoders on the MEG signal. Top Panel: Decoders with all Within community epochs (Familiar & New) vs. all Between communities epochs (Familiar & New) transitions. A) Generalization Across Time (GAT) matrix with significant cluster delineated in black. B) Time-by-time decoding. The shaded area indicates a significant temporal cluster. C. Individual performances based on whole epoch decoding: Mean Decoding accuracy across subjects (green bar, one dot per subject, the black line represents standard error). Those three analyses have been replicated with Familiar only transitions (middle panel) and New only transitions (bottom panel). Community structure was encoded in each case despite the flat local transition probability. Stars represent significance of the statistical tests (* p<0.05, ** p<0.01, ***p<0.001).

To assess robustness, we replicated the decoding accuracy with a different metric, we performed a decoding analysis on the whole epoch at the subject level for training and testing. This decoder simultaneously used all time points across all recording channels, providing a single accuracy value for the entire epoch. Unlike time-by-time decoding, this approach can exploit the temporal dynamic of the signal to differentiate conditions.

For the previous analysis, we pulled together the data from *Familiar* and *New* transitions. In a further analysis, we investigated whether the success of decoding the community remained possible when analyzing *Familiar* and *New* transitions separately. Therefore, we replicated the previous decoding analysis but limited it to *Familiar* transitions only, which had identical high local transition probabilities of 23% (*Familiar Within* Vs *Familiar Between*), or to *New* transitions only (*New Within* Vs *New Between*), which had a low transition probability of 4%. Note, however, that in this last case, the number of epochs was small, resulting in a low signal to noise ratio.

#### Statistical Analysis

Statistical significance in the Generalization Across Time (GAT) matrix was assessed using a temporal-temporal cluster-based permutation (MNE python ***(Gramfort et al., 2013)***) for times between 0 and 300ms. For the time-by-time decoder, we performed a temporal cluster permutation test in [0, 300]ms time window. Note that these two statistical tests are not independent as the time-by-time decoding corresponds to the diagonal of the GAT Matrix. The whole epoch decoding gives a single decoding value per subject, we thus performed a one-way t-test across subjects to test whether the decoding performance was significantly above-chance.

### Long-horizon associative learning estimation and linear regression

To assess the duration of the representation of a sequence item in the brain signals, we used epochs containing 10 tones (2.5s). We trained a 12-class decoder (for the 12 tones) with balanced accuracy to decode the identity of the first tone of the epoch throughout the whole epoch. To ensure that we were decoding the sustained activity related to the first tone and not a subsequent repetition of the same tone, we removed from the analysis all epochs in which the first tone was repeated during the test window (~ 65 % of the epochs were removed, fig 4A). We averaged the above chance decoding performance over the time-windows (250ms), which corresponded to the interval between two consecutive items, to estimate the amount of superposition of the representations of the different elements of the sequence. We then estimated the long-horizon associative learning strength of the association for each pair (Â), which corresponds to the sum of the transition probability matrix between the tones at all orders (A^t^), weighted by the overlap between item representations (fig 4B).

We later used the associated novelty index, defined as the negative log of this association strength, as a regressor for the MEG signal during the short epochs corresponding to the different transition types (fig 4D). We performed spatio-temporal cluster analysis on the beta value associated with this linear regression to extract electrodes and times where this long-horizon associative learning estimation might significantly explain the difference in activity across conditions. We also computed the average association strength of each type of transition (fig 4C).

## Results

Our experiment aimed to identify the neural correlates of community structure encoding and evaluate if this learning stems from a low-level associative process or corresponds to a late and explicit discovery ***(Ren et al., 2022; Stiso et al., 2022)***. To assess the encoding of the community structure, we first decoded *Within* vs *Between* transition type to characterize the temporal dynamics of the representation of each tone in the sequence in order to assess the possibility of overlapping representations that might allow long-distance associations. Based on this measured overlap, we could estimate the long-horizon associative familiarity for each transition. Finally, to determine whether this long-horizon associative learning model was indeed a plausible hypothesis, we ran a linear regression between the predicted familiarity and our data.

### Decoding Within Vs Between community transitions

We first tested whether participants’ mental model of the sequence encoded the community structure despite uniform transition probabilities. We thus trained and tested decoders on all tone epochs ending in a *Within* transition vs all tone epochs ending in a *Between* transition on all pairs (*Familiar* and *New)*. We obtained a significant cluster (p<0.05) in the GAT matrix accuracy. Temporal cluster analysis on the Time-by-Time decoding accuracy revealed a significant cluster between 90 and 250ms (p<0.001), peaking at 160ms. Finally, the epoch-based decoding was significantly above chance (p<0.01) (see fig 2 *Within* vs. *Between*).

We then restricted this analysis to the *Familiar* transitions (*Familiar Within* vs. *Familiar Between*, which corresponds to 92% of the epochs). Since *Familiar Within* and *Familiar Between* transitions have the same transition probabilities (0.23), a significant difference would then be due to a higher-order representation of the community structure. Here again, a significant cluster (p<0.01) was found in the GAT matrix. A temporal cluster between 80 and 280ms was found in the time-by-time decoding (p<0.001) with a peak at 150ms. Epoch-based decoding was also significantly above chance (p<0.001). Symmetrically, we restricted the analysis to *New* transitions only (*New Within* vs. *New Between*, which corresponds to 8% of the epochs). By design, both *New Within* and *New Between* transitions had transition probabilities of 4% so learning only local transition probabilities would predict equal unfamiliarity with both types of transition. In line with the previous results, we found a significant temporal-temporal cluster in the generalization matrix (p<0.05), and a significant temporal cluster in the time-by-time decoding (p<0.05, significant time = [130, 170] ms, peaking at 160ms). Epoch based decoding was also significant (p<0.05). Due to the much smaller number of epochs, the results were noisier.

We also computed the ERF on the gradiometers for the *Familiar* Within vs *Familiar Between* and *New Within* vs *New Between* contrasts on the [100-200]ms time-window to confirm the presence of the effect found with the decoding approach. The outcomes were qualitatively comparable: a significant effect around 150 ms for the *Familiar Within* vs. *Familiar Between* contrast (cluster-based permutations p<0.001), and a trend effect for the *New Within* vs. *New Between* contrast (cluster-based permutations p=0.075). In both cases, the topography of the difference was compatible with an auditory response.

We performed a series of control analyses to eliminate putative low-level confounds, such as decoding success based on the identity of the current tone, the previous tone, or the pair of tones. To control for tone identity decoding, we ran the decoding analysis but restricted it to one of the four nodes at the border of a community (i.e., connected to a node of the other community, darker nodes in fig 3). Depending on the previous tone, these epochs could be either *Familiar Within, Familiar Between, New Within*, or *New Between*. Thus, decoding within vs between community transitions on those epochs cannot be driven by the tone identity. The same was done for epochs where the transition began with one of these four nodes (i.e., epochs where the previous node of the sequence was one of the nodes at the border of communities) to control for decoding the identity of the previous tone. We also controlled for the pair forming the transition (previous and current tone identity simultaneously): in a similar manner to the current tone control, we restricted the analysis to nodes at the borders of communities and also cross-validated the decoding on the previous tone identity. To do so, we trained and tested our decoder on different previous nodes (training on three previous nodes per community and testing on the three others, see batches in fig 3). This strategy was also used for the *Familiar Within* Vs *Familiar Between* Generalization Across Time matrix (GAT) decoder. By experimental design, *New Within* vs. *New Between* decoders were already balanced for current and previous tones (each node is attached to one transition of each type). Thus, we only controlled for the pair by using the cross-validation of the previous node with the same batches described above. Overall, the control analyses qualitatively and quantitatively confirmed previous results. Only the *New Within* vs. *New Between* control for pair (i.e. controlling both previous and current tone identity simultaneously) analysis did not reach significance, probably due to the small number of epochs in this analysis (only 8% of the data was used in this last control).

**Figure 3:**
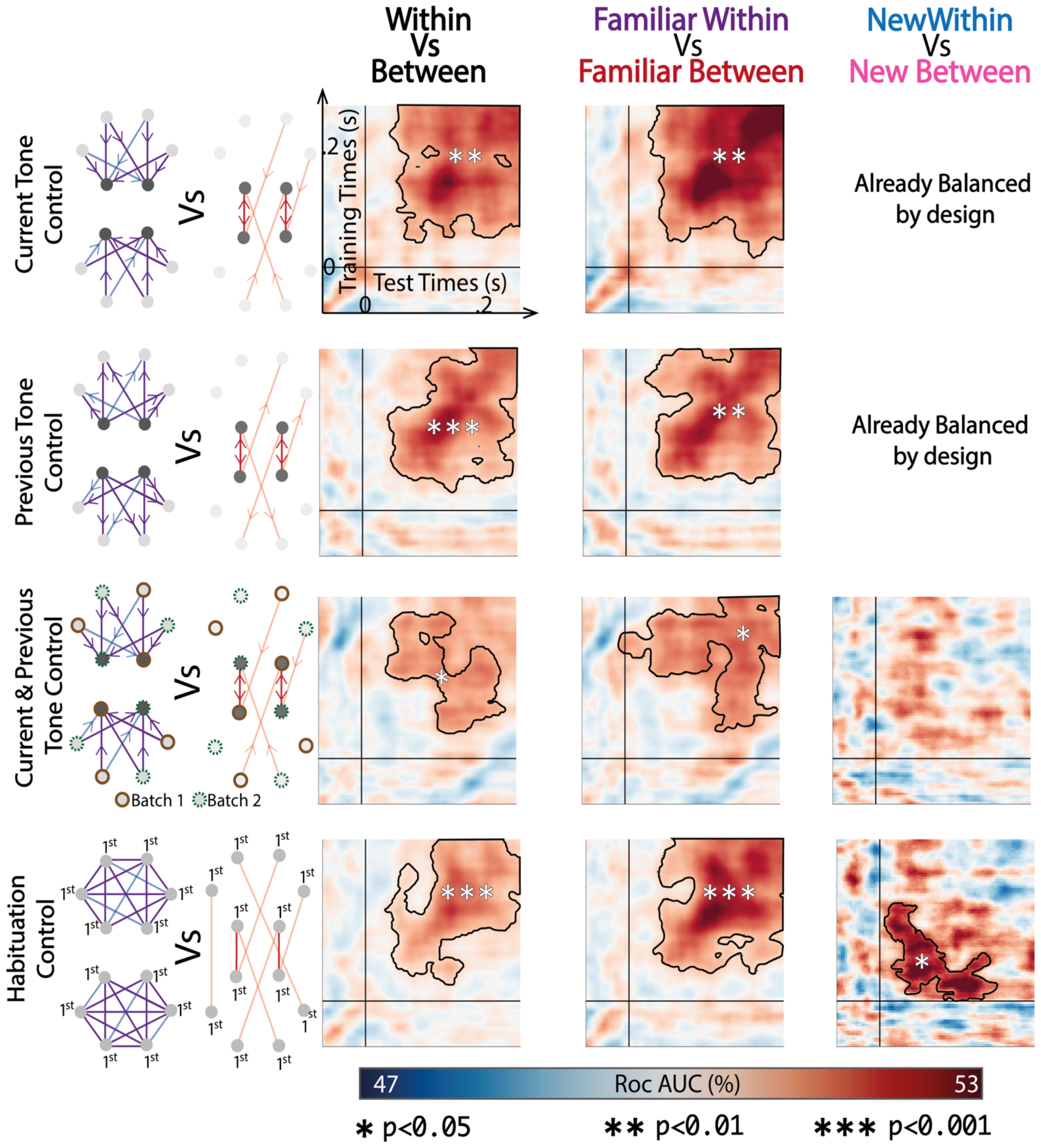
Control analyses for the results presented in figure 2. For each decoder, we controlled for the current tone, the previous tone and the pair (both current and previous tone simultaneously). We also controlled for habituation due to temporal proximity between tones. All the analyses qualitatively and quantitatively confirmed previous results except the New Within vs. New Between control analysis that did not reach significance, probably because of the small number of epochs.

*Within* vs *Between* community transitions decoding could also rely on a habituation effect. Indeed, if the sequence remains within a community, a particular sound might be repeated multiple times within a short span, causing habituation. However, if the sequence shifts from one community to another, the same sound is less likely to be repeated in a short time, thus preventing habituation. Therefore, this differential habituation effect could drive the *Within* vs. *Between* decoder. To rule out this alternative hypothesis, we restricted the analysis to the first appearance of each tone after a community change. Thus, close repetitions of tones of the same community are avoided in the data used for this decoder. Despite a decrease in the number of epochs, the decoding accuracy of those controls was still significant for all conditions. All generalization matrices are shown in fig 3.

### Long-horizon associative learning estimation

We tested here the hypothesis that long-horizon associative learning (associative learning over several consecutive and non-consecutive items) can support the encoding of network structure. This concept builds on Hebbs’ principle of strengthening the link between co-occurring events. Nonetheless, instead of focusing solely on learning adjacent pairs, we proposed a broader approach that allows connections to be established over longer distances. In our experiment, this long-horizon associative learning implies that the mental representation of each tone is sustained for a sufficient duration to allow several tones to overlap ***(Endress, 2010)*** and thus enable associations through more successive tones. According to this model, it is predicted that the representation of each tone should decrease following an exponential profile. To test this hypothesis, we quantified the overlap between the representations of item *n* and item *n+i*. In fact, this provides a good estimator of the weight of the non-adjacent transition probability of order i.

To estimate the overlap between brain representations of different items of the sequence, we determined how long the representation of each item was seen in brain activity. To do so, we split the data into 10-items long sequences (i.e. 2.5 seconds) with no repetition of the first tone in the sequence. We train a 12-class decoder on each time-point to predict the identity of the first tone. Decoding performance is shown in fig 4. We averaged the above chance decoding performance over the time-windows corresponding to the interval between two consecutive items. We observed an exponential-like decrease in performance that reached 0 after ~ 8 sequence items (fig 4A). It shows that the overlap enabling associative learning might thus include long-horizon dependencies of up to 8 items.

**Figure 4:**
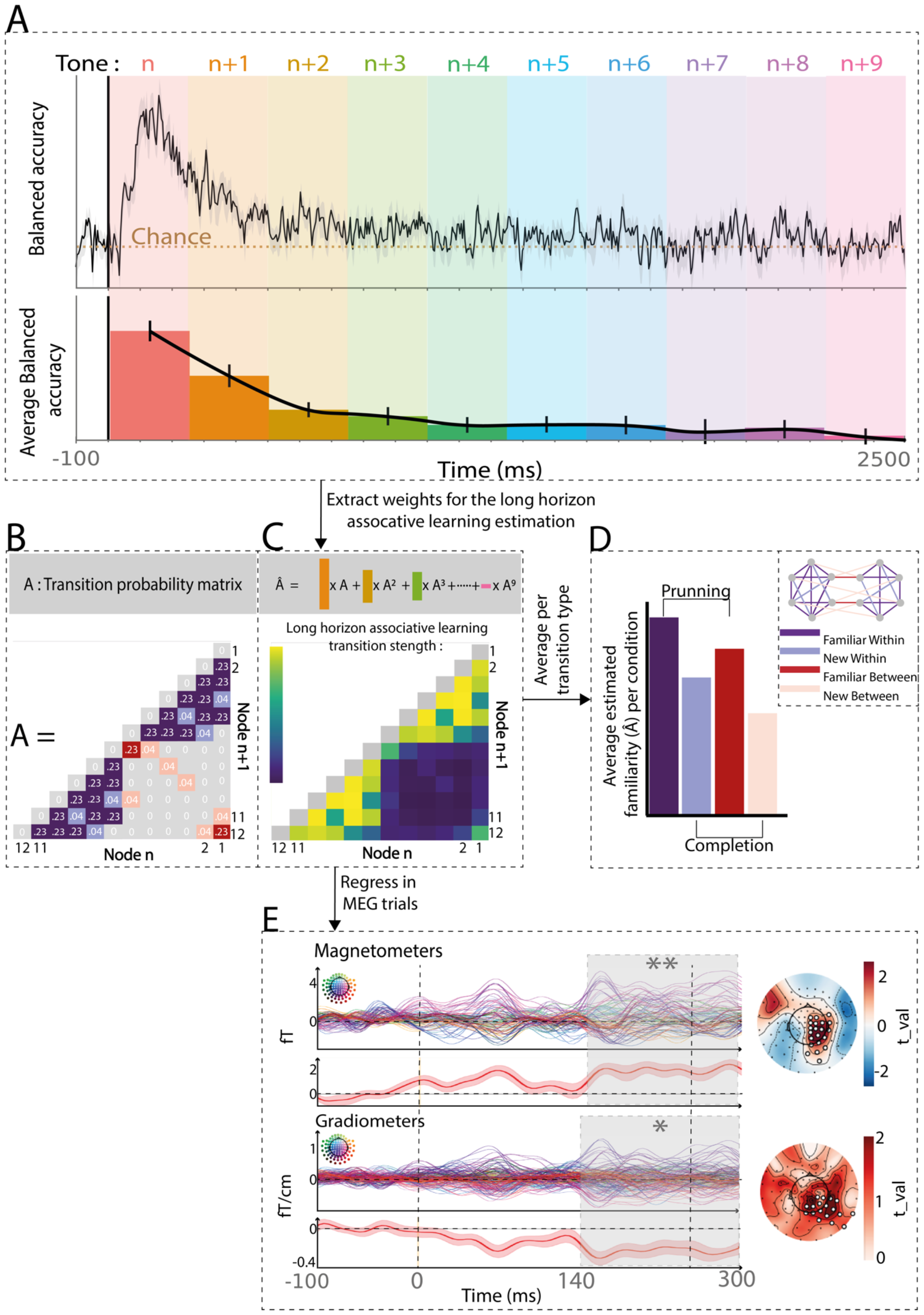
Associative learning estimation and fit on MEG data. A) Top: Decoding performance of the first item of the sequence across time (2.5s window). Shaded colors indicate the Stimulus Onset Asynchrony (SOA) between each tone of the sequence. The dotted line shows the chance level. Bottom: Decoding performance averaged over the duration of each tone and the following Inter Stimulus Interval (ISI). Error bars present the standard error across subjects. It takes ~8 items for the decoder of the first tone to converge to chance level. B) Matrix of exact transition probabilities (A) associated with the graph underlying the sequence. Familiar transitions are associated with 23% transition probabilities and New with 4% (fig1). Impossible transitions have a null transition probability. C) Estimation of the long-horizon associative learning strength for each transition. Based on the decoder (panel A), we estimated the overlap between non adjacent elements of the sequence (average decoder accuracy during SOA of item n+i). We then computed the associative learning strength (Â matrix) for each pair of elements as the sum of the different transitional probability orders (A^t^), weighted by the overlap between item representations. D) Average of the long-horizon associative learning strength per condition. Pruning (Familiar Within > Familiar Between) and completion (New Within > New Between) effects are consistent with behavioral results (Benjamin et al., 2023a)) and with the decoding performance obtained in fig 2. E) Regression coefficient for the estimated long horizon associative novelty (-log(Â)) for each MEG sensor. Significant time-windows are shown in shaded areas and significant sensors are indicated on the t-map topographies by the white dots. These were obtained with a spatio-temporal cluster-based permutation analysis. The red line below the sensors value represents the time course of the average regression value on the sensors of the significant cluster.

We estimated the long-horizon associative learning strength of each pair of tones. To do so, we computed the sum of the different transitional probability orders weighted by the overlap between item representations as estimated from the decoding performances (fig 4B). This gave us a 12x12 symmetrical matrix of learning familiarity for each pair (fig 4C). Finally, we averaged this measure of Familiarity for each condition type (fig 4C) and obtained a result that is consistent with the pruning effect (difference between *Familiar Within* vs *Familiar Between* transitions) and the completion effect (difference between *New Within* and *New Between* transitions) as discussed in ***(Benjamin et al., 2023a)***.

### Long-horizon associative learning accounts for epoch variability

To test the neural predictions of long-horizon associative learning, we correlated brain signals with the estimated associative learning strength of each transition (fig 4D). We performed a linear regression between the brain signal after each tone and the novelty effect produced by each transition. Unlike most studies of sequence learning, where the novelty is calculated solely from local transition probabilities, we computed it here as the negative log of the long-horizon associative learning strength. This calculation takes into account several orders of adjacent and non-adjacent transition probabilities whose weights have been computed based on the overlap of brain representations estimated by our tone decoder (fig 4 A-D). A spatio-temporal cluster permutation test revealed a significant cluster (fig 4E) in the magnetometers (right centro-occipital, time = [150 ; 290] ms, pval < 0.01) that was replicated in the gradiometers (right centro-occipital, time = [140 ; 300] ms, pval < 0.05).

Furthermore, the observed clusters were still significant when the negative log of the adjacent transition probabilities was introduced as a supplementary regressor (ps < 0.05 for both magnetometers and gradiometers clusters). However, if the high correlation between the TP matrix and the long horizon-associative model makes it hard to directly disentangle those two models solely based on this regression analysis, it does nicely complement the decoding analyses.

## Discussion

In this study, our aim was to determine whether local statistical learning and structure learning in sequences are governed by the same cognitive process or by distinct processes. Learning local statistics is often described as an associative process, while network learning is usually seen as an abstract map representation. Previous studies exploring network learning have used explicit paradigms, revealing late brain signatures consistent with top-down or frontal activity ***(Ren et al., 2022; Stiso et al., 2022)***. However, based on a modeling approach, we proposed in our previous behavioral study that low-level associative learning strategies might support both local and high-order statistical scales ***(Benjamin et al., 2023a)***. Thus, this hypothesis predicts that learning sequence structure does not require an explicit representation and may instead rely on automatic and rapid (~150ms) mismatch responses, similar to those observed after the violation of local transition probabilities.

### Network learning results from a low-level bottom-up computations

To test these predictions, we presented participants with a passive learning task using rapid auditory sequences. We showed that the structure properties of the sequence were rapidly decodable from the participants’ brain recordings (~[100-250] ms after tone onset). The timing of this response, as early as 150ms after the information became available, aligns with the rapid deviant responses (MMN in EEG) observed in learning based on violation of transitional probabilities ***(Maheu et al., 2019; Todorovic and de Lange, 2012)***. Since the transition probabilities between tones were uniform and the walk within the network was random, prediction could not be based on high-level top-down expectation. This early and automatic response (150 ms after the transition) challenges the notion of abstract and explicit calculations as prerequisites for learning such structures. In addition, our analyses revealed a similar effect when the decoding analysis was restricted to new transitions (*New Within* Vs *New Between*) and to familiar transitions (*Familiar Within* Vs *Familiar Between)*, suggesting an automatic generalization of the community structure beyond sensory evidence. This result provides a neural underpinning for the behavioral observations we previously reported, indicating that participants accurately assess the familiarity of transitions based on their congruence with network structure, even when these transitions were not encountered during training.

### Long-horizon associative learning as a plausible implementation for FEMM

In our previous study, we hypothesized that the FEMM could effectively explain adult behavioral performance. This model aggregates the different orders of statistical regularities (adjacent and non-adjacent) into a single quantity. In this study, we showed that this model can be readily implemented through a simple associative learning mechanism relying on Hebb’s principle ***(Benjamin et al., 2023a; Hebb, 1949)***. In the context of structure learning, this principle would imply a sustained mental representation of each tone for a sufficient duration to enable the overlapping of several elements despite the temporal distance. We thus predicted the representation of each tone to exhibit an exponential decay profile. A rapid decay of tone information would limit associations to short distances, while a slower decay would facilitate the formation of long-horizon dependencies and, therefore, the extraction of the underlying structure. Thus, this exponential decay acts as a balance between local relevance and generalization.

To test this idea, we estimated the duration of the representation of each tone, performing a decoding analysis of tone identity. The identity of a tone was decodable during the presentation of the subsequent eight tones, with a decoding performance exponentially decaying over subsequent tones. This profile provided an estimation of the number of elements simultaneously represented at a given time. Consequently, it allowed us to quantitatively assess the strength of each tone pair in the heard sequence (fig4C). We found that these weights accurately accounted for the results of the *Within* vs. *Between* decoders, encompassing both *Familiar* and *New* transitions (Fig. 4D). Moreover, this estimated strength significantly correlated with neural activity, aligning with the timing of the automatic deviant response ***(Maheu et al., 2019; Todorovic and de Lange, 2012)***. This result provides compelling evidence for the rapid encoding of structure through bottom-up processes compatible with associative learning strategies.

However, it is worth noting that an alternative implementation of the same metric is theoretically possible. Simple pairwise association learning, in combination with a transitivity property, would also predict similar learning. In fact, if participants solely learn pairs (e.g., A-C and C-D), transitivity of this learning can strengthen the A-D pair, even if not explicitly presented. Considering this transitivity with similar exponentially decreasing weights would be mathematically equivalent to our model while not strictly requiring a sequential presentation of the structure. Although, we cannot definitively rule out this alternative implementation of the same metric, our findings suggest that sequential presentation is crucial to have an overlap between successive items representations, enabeling Hebbian associative learning., It is also important to acknowledge that associative learning might not be the sole mechanism contributing to network structure learning, particularly in cases where explicit detection is required from participants. Abstract representations of hippocampal maps ***(Constantinescu et al., 2016)*** or frontal maps ***(Stiso et al., 2022)*** might also play a role in such tasks ***(Garvert et al., 2017; Schapiro et al., 2017, 2016)***. Intracranial recordings conducted during local statistical learning paradigms have revealed that multiple brain regions, including cortical areas and hippocampus, can simultaneously represent the same structure while carrying different information ***(Henin et al., 2021)***.

### Difference between implicit passive listening and explicit structure learning

Thus, converging results provide evidence that associative learning supports the perception of the community structure in the present experiment. Long-horizon associative learning strength significantly accounted for the variance in brain signals (fig 4E). Moreover, the pruning and completion effects found with decoders (fig 2) can easily be explained by the same mechanism (see fig 4D). However, it is worth noting that the results from our previous behavioral study do not entirely align with the current ones. Specifically, in the present experiment, the representation of tones exhibited a more rapid decrease (exponential decrease factor 0.52) as compared to its estimation in our previous behavioral study (factor 0.058, about ten times lower). This discrepancy suggests that participants in the current experiment might be less inclined to generalize the underlying structure.

Several factors might explain this difference. Firstly, the generalization factor estimation in the MEG experiment may be noisier due to the small number of participants (23 vs. several hundred in the behavioral study). Since the trade-off between generalization and accuracy may vary among individuals ***(Lynn et al., 2020)***, on one side group level estimation with 23 subjects is limited and on the other side at the individual level, it is difficult to measure this trade-off due to the data variability. A larger sample size with multiple sessions per subject would be necessary to obtain a reliable estimation of the generalization factor at the individual level. Secondly, it is possible that associative learning represents the implicit component of this task ***(Andringa and Rebuschat, 2015)***, followed subsequently by an explicit decision-making process involving higher level prefrontal regions. This second step might facilitate the abstraction of the structure by labeling each community as distinct ***(Koechlin et al., 2003; Koechlin and Jubault, 2006)***. This dual process could explain why explicit behavioral tasks ***(Benjamin et al., 2023a; Lynn et al., 2020)*** exhibit a better generalization factor compared to our implicit MEG task. The same explanation may account for the late signatures of top-down activity reported by ***(Ren et al., 2022)*** who used a slow and explicit task. To further explore this hypothesis, a direct comparison of passive and active learning of such networks while monitoring the representations in the auditory cortex, the hippocampus, and the lateral prefrontal cortex would be necessary.

## Conclusion

The aim of the present study was to uncover the neural mechanism underlying network learning. We proposed the sparse community paradigm as a way of combining local statistical learning and network learning in a single sequence. Previous behavioral studies have shown that a mathematical model (FEMM) accurately captures human learning. Here, we add that the behavioral pattern described by the FEMM is compatible with certain associative learning principles. Indeed, thanks to time-by-time decoding of the brain state associated with a tone, we observed an exponential decay in the tone representation across 8 elements. Using this estimate of mental representations’ dynamics, we estimated the strength of each network transition. This estimate significantly correlated with our data. The present study provides novel insights into the mechanism underlying network learning and highlights the importance of brain dynamics in the understanding of sequence learning. Further investigations in different experimental conditions (explicit vs implicit), over different tone and ISI durations, with different populations (non-human primates), and during early development are necessary to better characterize this learning ability.

